# Gamma oscillations in the rat ventral striatum originate in the piriform cortex

**DOI:** 10.1101/127126

**Authors:** James E. Carmichael, Jimmie M. Gmaz, Matthijs A. A. van der Meer

**Author notes:** Correspondence should be addressed to MvdM, Department of Psychological and Brain Sciences, Dartmouth College, 3 Maynard St, Hanover, NH 03755.

## Abstract

Local field potentials (LFP) recorded from the human and rodent ventral striatum (vStr) exhibit prominent, behaviorally relevant gamma-band oscillations. These oscillations are related to local spiking activity and transiently synchronize with anatomically related areas, suggesting a possible role in organizing vStr activity. However, the origin of vStr gamma is unknown. We recorded vStr gamma oscillations across a 1.4mm^2^ grid spanned by 64 recording electrodes as rats rested and foraged for rewards, revealing a highly consistent power gradient originating in the adjacent piriform cortex. Phase differences across the vStr were consistently small (<10^°^) and current source density analysis further confirmed the absence of local sink-source pairs in the vStr. Reversible occlusions of the ipsilateral (but not contralateral) nostril, known to abolish gamma oscillations in the piriform cortex, strongly reduced vStr gamma power and the occurrence of transient gamma-band events. These results imply that local circuitry is not a major contributor to gamma oscillations in the vStr LFP, and that piriform cortex is an important driver of gamma-band oscillations in the vStr and associated limbic areas.

**Significance Statement:** The ventral striatum is an area of anatomical convergence in circuits underlying motivated behavior, but it remains unclear how its inputs from different sources interact. One of the major proposals of how neural circuits may dynamically switch between convergent inputs is through temporal organization reflected in local field potential (LFP) oscillations. Our results show that in the rat, the mechanisms controlling vStr gamma oscillations are primarily located in the in the adjacent piriform cortex, rather than vStr itself. This provides a novel interpretation of previous rodent work on gamma oscillations in the vStr and related circuits, and an important consideration for future work seeking to use oscillations in these areas as biomarkers in rodent models of human behavioral and neurological disorders.

## Introduction

The ventral striatum (vStr) is an anatomically heterogeneous region receiving a convergence of anatomical connections from structures in the cortico-striatal-thalamic loop, as well as from the hippocampal formation and amygdala (Pennartz et al., 1994; Haber, 2009; Sesack and Grace, 2010). Prominent local field potential (LFP) oscillations can be recorded from the rodent and human vStr, spanning a broad range of frequencies that include delta, theta, beta, multiple gamma bands, and high-frequency oscillations (Berke et al., 2004; van der Meer et al., 2010; Axmacher et al., 2010; Dürschmid et al., 2013; Hunt et al., 2010) and display phase-amplitude coupling (humans: Cohen et al. 2009a, rodents: Donnelly et al. 2014; Malhotra et al. 2015). A particularly salient feature of the vStr LFP is the presence of prominent gamma-band oscillations, which include distinct low-gamma (∼45-65 Hz) and high-gamma (∼70-90 Hz) components.

Several studies have found correlations between the occurrence of these low- and high-gamma oscillations such as reward approach and/or receipt (van der Meer and Redish 2009; Berke 2009, but see Malhotra et al. 2015), reward expectation and prediction errors (Cohen et al., 2009a; Axmacher et al., 2010), drug-related conditioned place preference (Dejean et al., 2016), and impulsive actions (Donnelly et al., 2014). vStr gamma oscillations are also affected by manipulations of the dopamine and cannabinoid systems (Berke, 2009; Lemaire et al., 2012; Morra et al., 2012). Importantly, the spiking of vStr neurons is related to these gamma oscillations, with putative fast-spiking interneurons (FSIs) displaying a particularly striking pattern: partially distinct populations appear to phase-lock to low- and high-gamma respectively (Berke, 2004; van der Meer and Redish, 2009; Dejean et al., 2016). Medium spiny neurons (MSNs), which make up the vast majority of vStr neurons, tend to show weaker but nonetheless statistically significant phase locking (Kalenscher et al., 2010; Howe et al., 2011) and ensemble spiking activity can predict which gamma band oscillation is present in the LFP (Catanese et al., 2016).

A conservative interpretation of the above body of work is to view gamma oscillations in the vStr LFP as a signal containing a certain amount of information about the state of the local network, that is, as a practically useful readout. However, in several brain structures there is evidence that the effectiveness of an incoming stimulus can depend on the phase of local gamma oscillations (Cardin et al., 2009; Sohal et al., 2009); although such state-dependence has not yet been shown in the vStr, the findings reviewed so far suggest the possibility that the mechanisms generating vStr gamma contribute to dynamic gain control, as in “communication through coherence” proposals (Akam and Kullmann, 2010; Womelsdorf et al., 2014; Fries, 2015). This idea is conceptually attractive given the anatomical convergence of multiple inputs onto the vStr, which all oscillate (O’Donnell and Grace, 1995; Gruber et al., 2009; Harris and Gordon, 2015); indeed, gamma oscillations in the vStr LFP are often coherent with gamma-band LFP signals in anatomically related areas, with prefrontal cortical areas being the best-studied example (rodent: Berke 2009; McCracken and Grace 2009; Dejean et al. 2013;Catanese et al. 2016, human: Cohen et al. 2009b).

For both the “readout” or “gain control mechanism” views of vStr gamma oscillations, it is important to establish how this signal is generated; a crucial first step is to localize their source(s), which have so far remained unclear. Some studies emphasize the similarity of vStr gamma oscillations with those recorded in the adjacent piriform cortex (Berke, 2009) while other studies report local heterogeneities that appear to be inconsistent with volume conduction (Kalenscher et al., 2010; Morra et al., 2012), or focus on cell-intrinsic contributions such as gamma-band resonance (Taverna et al., 2007). Resolving this issue would help determine if local vStr circuitry may implement a “switchboard” through a communication-through-coherence type mechanism (Fries, 2005; Gruber et al., 2009; Akam and Kullmann, 2010), and to enable a productive dialog between rodent and human studies in which vStr LFPs have behavioral and clinical relevance (Sturm et al., 2003; McCracken and Grace, 2009; Dejean et al., 2013).

Thus, to determine the origin of gamma oscillations in the local field potential of the rat vStr, we (1) record from across the vStr using a high density electrode array, and (2) inactivate the piriform cortex using reversible naris (nostril) occlusions known to abolish piriform gamma (Zibrowski and Vanderwolf, 1997).

## Methods

### Overview

This study consists of two experiments: (1) a “LFP mapping” experiment measuring the distribution of gamma oscillations across the vStr, and (2) a “naris occlusion” experiment testing the effects of unilateral nostril closures on vStr gamma. Data in the LFP mapping experiment were acquired as rats performed a maze-based foraging task, as well as during off-task resting periods. Since we found no difference in the properties of gamma oscillations on- and off-task, data for the naris experiment were acquired during rest only. All procedures were approved by the University of Waterloo Animal Care Committee (AUPP 11-06) and the Dartmouth College IACUC (vand.ma.2).

### Subjects and timeline

Seven Long-Evans male rats (> 10 weeks old; >400g) were used in total (four in the LFP mapping experiment and three in the naris occlusion experiment). For the LFP mapping experiment, rats were pretrained on a foraging task (4 days of maze habituation), implanted with recording probes (described in detail below), retrained on the task following recovery (minimum 4 days), and recording data acquired. The naris experiment included two naive rats and a final rat that had been previously trained on set-shifting task used in Gmaz et al. (SfN abstract, 2016; no differences were found across rats). All animals were kept on a 12hr light/dark cycle with all the experiments performed during the light phase.

### Surgery and recording probes

Rats for the LFP mapping experiment were implanted with 64 channel silicon probes (A8×8-10mm-200-200-177, NeuroNexus; 50*μ*m thick). Probe recording sites and tetrodes were gold-plated (Sifco 6355) to impedances between 300-500kΩ (Nano-Z, White Matter LLC). Probes were attached to a microdrive and implanted as in Vandecasteele et al. (2012), with the addition of a separate independently movable stainless steel wire reference electrode (gold-plated to 50kΩ) implanted into the same hemisphere (AP 2.2 mm anterior to bregma, ML 2.0 mm, targeting the corpus callosum overlying the vStr). The probes contained regularly spaced recording electrodes, spanning 1.4mm^2^ and arranged in a 8×8 grid, and were implanted rotated around the vertical axis with the edges between AP 0.6–2.28, ML 1.4–2.8) (Figure 1C). Rats for the naris occlusion experiment were implanted with a either a custom built 4-tetrode multi-site drive, with one tetrode located in the vStr (AP 1.5, ML 2.0), with additional tetrodes in the orbitofrontal, prelimbic, and cingulate cortices (not analyzed here), or a custom 16-tetrode “hyperdrive” with all tetrodes in the vStr. Following surgery, probes and tetrodes were moved down over the course of several days until they reached the target region (DV 6.5–8 mm).

**Figure 1:**
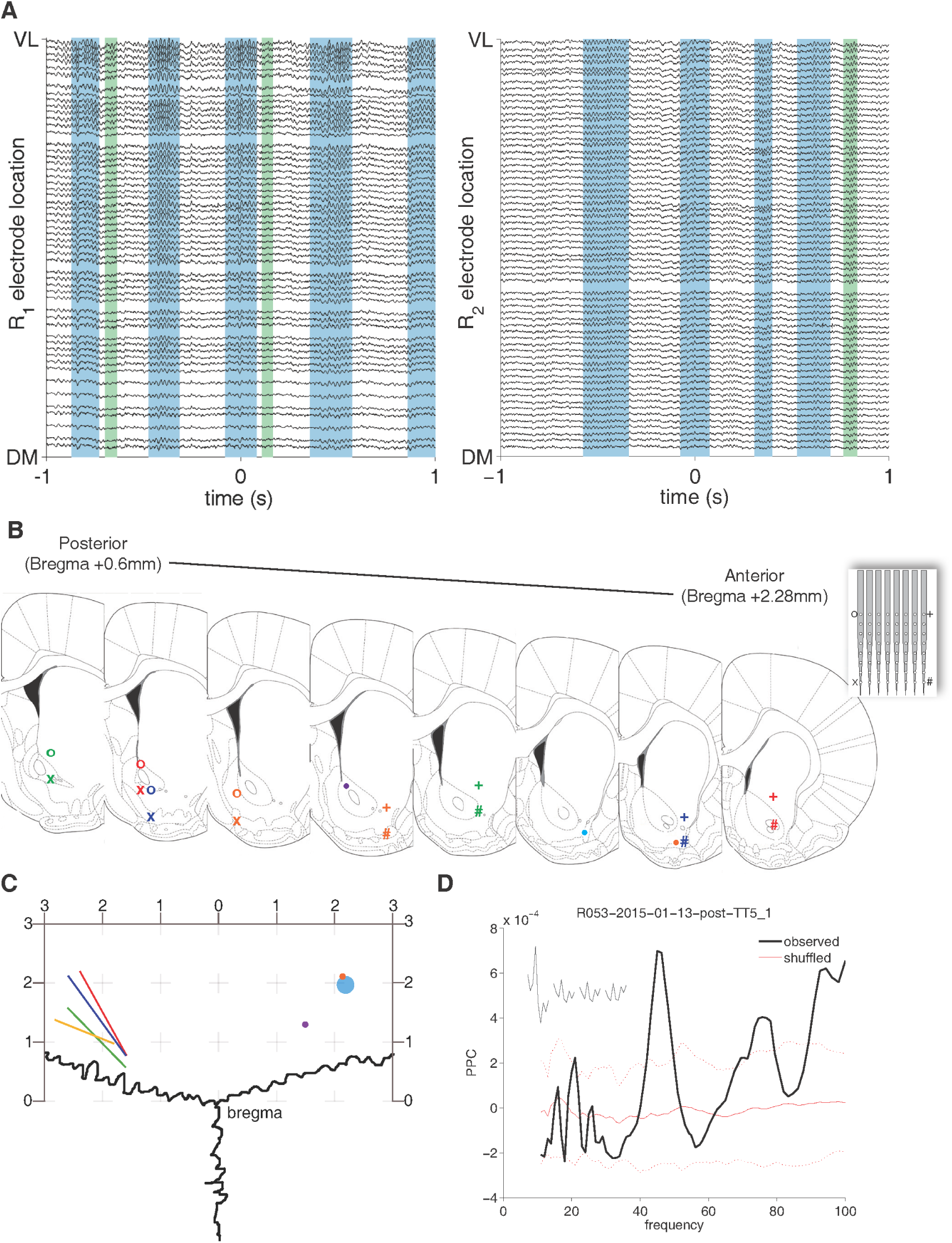
High-density planar silicon probe recordings across the ventral striatum. **A**: Example ventral striatum LFP traces (filtered between 1-500 Hz; the two sets are from different subjects R1 and R2) ordered by the location of the corresponding electrode site along the ventrolateral (VL, top) to dorsomedial (DM, bottom) axis of the probe. Gamma events were detected using a thresholding procedure; example low-gamma events are shaded in blue, high-gamma events in green. Note that the amplitude of the gamma events increases from the dorsomedial to ventrolateral electrodes. **B**: Recording locations as determined by histological processing. For silicon probes (inset), recording locations are indicated by the corners of the probe (o+x#), while tetrode locations are marked with dots (colored by subject) **C**: Dorsal view of the electrode locations to demonstrate the angle of the probes (lines), and location of tetrodes (dots) all colored by subject. **D**: Field pairwise phase consistency (PPC,Vinck et al. 2011) of a representative ventral striatal unit showing a clear peak in the gamma band.

### Behavioral task

The *LFP mapping* rats were trained on a square-shaped elevated track (width 10 cm, each edge of the square 100cm long) with sucrose reward receptacles placed at each of the four corners. Nosepoking in the reward receptacles yielded 0.1, 0, 0.1, or 0.2ml of 12% sucrose reward respectively; with experience (daily 40-minute sessions, preceded and followed by 5-10 minutes of off-task recording on a terracotta planter filled with towels) animals learn to skip the non-rewarded site. Throughout behavioral training and recording on the task, rats were food-restricted to >85% of their free-feeding weight to encourage foraging behavior. The *naris occlusion* rats had *ad libitum* access to food. Before the start of the experiments described here, one of the three animals had been previously trained on a behavioral task that used the same physical setup in the same room (Gmaz et al. SfN abstract, 2016).

### Data acquisition and preprocessing

Wideband signals were acquired for the silicon probe and “hyperdrive” recordings using RHA2132 v0810 multiplexing headstages (Intan) and a KJE-1001/KJD-1000 amplification system (Amplipex), sampled at 20kHz (decimated to 2kHz during analysis) referenced against a single electrode in the corpus callosum above the vStr. To extract spikes from the continuously sampled Amplipex data, the data was filtered (600-9000Hz), thresholded and peak-aligned (UltraMegaSort2k,Hill et al. 2011). Recordings for the naive multisite 4-tetrode drive rats used a Digital Lynx data acquisition system with an HS-18mm preamplifier (Neuralynx) with the low pass 0.5-500 Hz subsampled from 32kHz to 2KHz. Spiking data the voltage was recorded at 32kHz when the voltage exceeded a predefined threshold. Data for LFP mapping analysis had the DC offset removed and was filtered (1-500Hz, 10th order Butter-worth, `filtfilt.m`). Putative single neurons were manually sorted using MClust3.5 (A.D. Redish et al.). Electrode sites with irregular impedance values (>900kΩ), intermittent signals, or known defective sites (B-stock probes) were excluded from analysis (black pixels in Figures 2-5).

### Naris occlusion

Reversible naris closures (“nose plugs”) were constructed from PE90/100 tubing (Intramedic) by tying a human hair and a suture in a double knot, threading it through the side of the tubing, and gluing it to the inside of the tube such that it protruded a ∼8mm beyond the tubing, facilitating subsequent removal (Kucharski and Hall, 1987; Cummings et al., 1997). Nose plugs were coated with Vaseline and inserted (or removed) while the rat was briefly under isoflurane anesthesia (≤2 min from the time of induction). The effectiveness of this procedure in blocking air flow through the occluded nostril was verified by visual inspection: any remaining airflow tended to move the Vaseline. Daily recording sessions consisted of four off-task segments: a non-occlusion baseline (“pre”), ipsilateral and contralateral naris occlusions (order counterbalanced across sessions) and another non-occlusion baseline (“post”) separated by 45 minutes to minimize any effects of the isoflurane anesthesia (Figure 7G).

### Session inclusion criteria

For each *LFP mapping* subject, data from three consecutive daily recording sessions were analyzed, with the recording probe at the same depth across days. These sessions were chosen such that the probe had to be at the same depth across sessions, and that either the animal had reached the performance criterion on the task, and if they did not (2/4 rats) then the final three sessions at a consistent depth were used. For one LFP mapping subject, only two sessions were included due to a faulty recording tether. For the naris occlusion experiment, four days of data were recorded and analyzed (two days, followed by one day of no recording, and two more days).

### Data analysis overview

All analyses were performed using MATLAB 2014a and can be reproduced using code available on our public GitHub repository (http://github.com/vandermeerlab/papers) with data files, metadata, and code usage guides available upon request. We performed spectral analysis on the LFP data using power spectral densities (PSDs) over specific epochs within recording sessions (off-task baseline recordings before and after behavior; reward site approach (run) and reward site consumption; and rest data only for the naris occlusion group). Given that gamma oscillations in the vStr LFP tend to occur in distinct events (Masimore et al., 2005; Cohen et al., 2009b; Donnelly et al., 2014) we also performed event-based analysis (detailed below).

### Gamma event detection and analysis

Gamma events were detected following the procedure in Catanese et al. 2016. First, LFP data were filtered into the bands of interest (low-gamma: 45-65 Hz; high-gamma: 70-90 Hz) using a 5^th^ order Chebyshev filter (ripple dB 0.5) with a zero-phase digital filter (MATLAB `filtfilt`()). The instantaneous amplitude was extracted from the filtered signal by Hilbert transform (MATLAB abs(`hilbert`())). For the LFP mapping experiment, a single recording channel for gamma event detection was selected by first computing the power spectral density (PSD) for the six most ventrolateral recording sites and using the site with the largest power averaged across frequencies in the low-gamma range (45-65Hz). Large amplitude artifacts (>4 SDs from the mean in the unfiltered data) and chewing artifacts (>3SD of the z-scored data filtered in the 200-500Hz) were first removed from the data. If the instantaneous amplitude surpassed the 95^th^ percentile of the amplitude of the session in it was considered a candidate event. Events were excluded if they: co-occurred with high voltage spindles (>4SD in 7-11Hz), had <4 cycles, or had a variance score (variance/mean of cycle peaks and troughs) >1.5. Applying a threshold (40*μ*V) to the minimum filtered amplitude helped remove false positives in the high-gamma band only, but did not aid in rejecting false positives for low-gamma. script for both the mapping and naris data.

For each detected gamma event, a length-matched pseudo-random “control” event was extracted by identifying the period of lowest amplitude in the gamma band of interest within a 20s window prior to each detected gamma event. Gamma and random events were then converted into one of two formats. For gamma power analysis, events were converted to the FieldTrip format by taking 100ms of data centered on detected gamma events (`ft_redefinetrial`, FieldTrip toolbox,Oostenveld et al. 2011). For phase and current source density (CSD) analysis the events were converted into a three cycle “triplet” by identifying the cycle with the highest amplitude and extracting it as well as the cycle before and after. The three extracted cycles were then interpolated to ensure that each event had an equal number of samples (214) allowing for averaging across events.

For the naris occlusion experiment events were detected using the percentile method detailed above, except that a threshold value (in microvolts) was obtained from the baseline data and then applied to the ipsi- and contralateral sessions; this was necessary to be able to compare the number of gamma events between conditions.

### Spectral analysis

Session-wide PSDs were computed for the naris experiment using Welch’s method (∼2s Hanning window) on the first derivative of the data to remove the overall 1/f trend (MATLAB `pwelch`(`diff`(`data`))). Event-based gamma power was computed using the FieldTrip toolbox (Oostenveld et al., 2011) function `ft_freqanalysis` (’mtmfft’, Hanning window, frequency of interest (FOI) 1-500Hz, window 5./FOI) by averaging the frequencies of interest in the resulting PSD. Phase differences were computed relative to the ventrolateral most electrode using cross-power spectral density (MATLAB `cpsd`) using a 64 sample Hamming window with 50% overlap (NFFT = 1024).

### Plane fits

To quantify the consistency of any patterns in the gamma power distribution across probe recording sites we computed a plane of best fit using least squares for the gamma power across the array during each gamma event and compared the variance explained (R^2^) to the pseudo-random non-gamma events of equivalent length.

### Current source density analysis

Current source densities (CSD) were computed by taking the second spatial derivative of the bandpass filtered data across the recording channels along the dorsomedial to ventrolateral diagonal of the recording array and multiplying it by the conductance (0.3mS/mm). Missing channels along this diagonal were filled in by interpolation (MATLAB `griddata`). Average CSDs for each rat were computed using the three cycle triplet from each detected gamma event (mentioned above).

## Results

### Experiment 1: mapping of gamma oscillations in the ventral striatum

Clear gamma events were recorded across the vStr (Figure 1A) of all rats chronically implanted (n = 4) with 64-channel planar silicon probes, which covered an area of 1.4mm^2^ with a regular 8×8 grid of recording sites (Figure 1B and C). We recorded wideband neural data during behavior on an elevated maze and during off-task resting periods. As in previous reports, we found clear low- and high-gamma oscillations in the LFP (Figure 1A; Leung and Yim 1993;Berke et al. 2004;Howe et al. 2011) using probes implanted in the vStr, to which putative interneurons showed significant phase locking (Figure 1D). Gamma oscillations appeared highly coherent across sites, with visual inspection suggesting systematic changes in power across sites. To quantify this effect, we isolated gamma events using a thresholding procedure on the channel with the largest average gamma-band power (*Methods*); examples of detected events are shown as shaded areas in Figure 1A (low-gamma: blue, high-gamma, green). We plotted the gamma-band power across all probe sites as a heat map, illustrated for an example low-gamma event in Figure 2A and B. An approximately linear gradient was apparent, with the highest power at the ventrolateral pole and the lowest power at the dorsomedial pole (Figure 2B).

**Figure 2:**
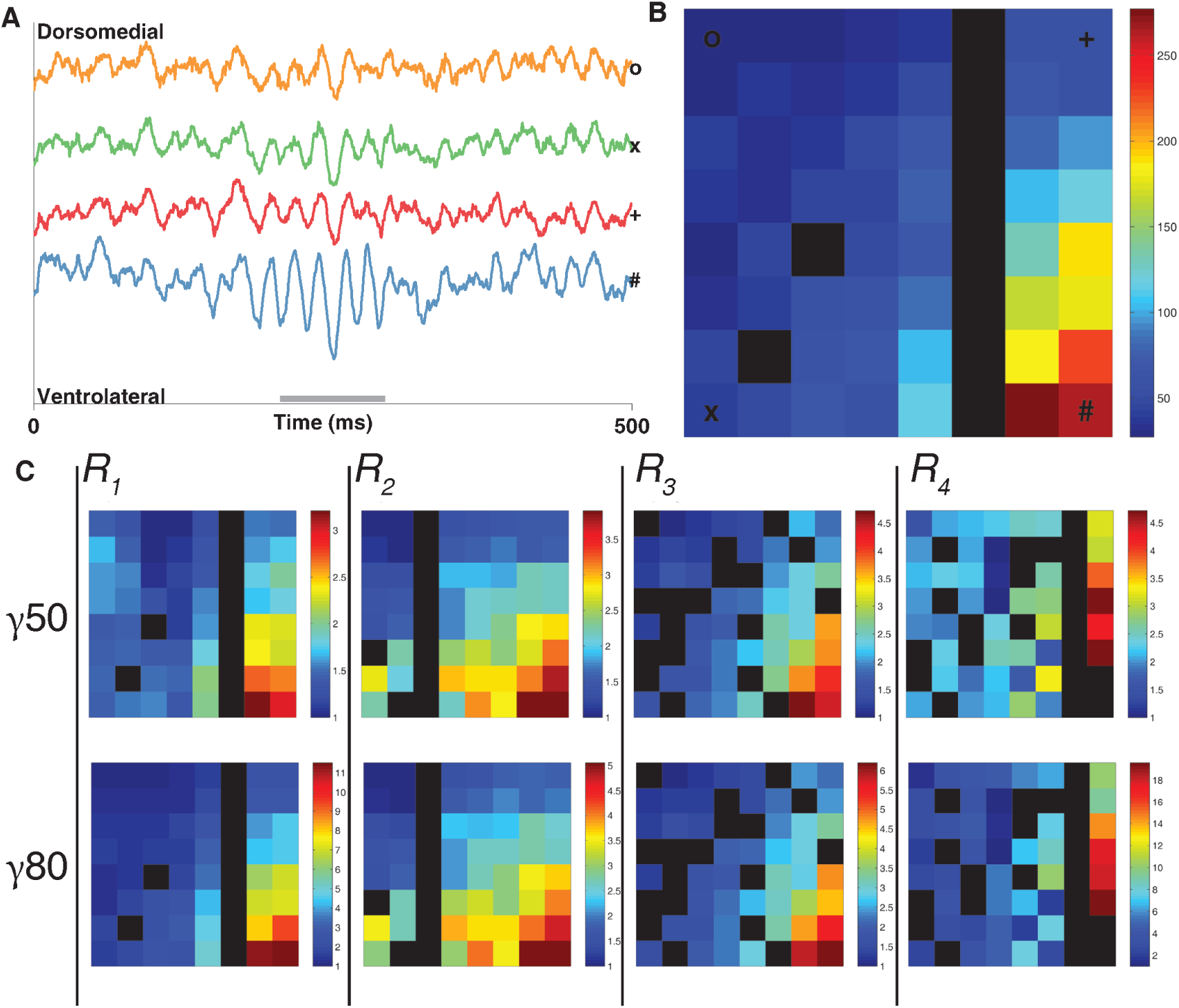
Gamma band local field potential (LFP) oscillations in the ventral striatum form a dor-somedial to ventrolateral power gradient. A: Raw LFP traces (left) from the recording electrodes at the dorsomedial (o), ventromedial (x), dorsolateral (*) and ventrolateral (#) points of the silicon probe for a representative low-gamma event. Grey bar spans the length of the gamma event (see *Methods* for details of gamma detection). B Heat map (right) showing the gamma power (*μV*^2^) across the recording array (64 sites, regularly spaced in an 8×8 grid spanning 1.4mm^2^) during the same low-gamma event as seen in the raw traces (left). Gamma power is about five times greater in the ventrolateral region compared to the dorsal-medial region. C: Average low- and high-gamma power across an entire recording session for each subject using the same probe layout in similar recording locations across animals (scale is normalized to the lowest power channel). Black spaces represent defective recording sites (see *Methods* for defective site criteria).

As a first step towards establishing the generality of this linear gamma power gradient, we detected low-gamma (45-65 Hz) and high-gamma (70-90 Hz) events across multiple recording sessions and subjects (11 sessions across 4 subjects; total events detected, 5997 for low-gamma, 5005 for high-gamma; mean event rates 0.32/s for low-gamma, 0.32/s for high-gamma; mean ± SD event length, 140.89 ± 59.52 ms for low-gamma, 81.30 ± 42.69 ms for high-gamma). Although the distribution of functional sites varied between probes, the linear gamma power gradient was consistent across subjects (Figure 2C). Next, we asked if the distribution of gamma power across the vStr was affected by activity level (running on an elevated track for sucrose reward vs. off-task rest) or by different behaviors during the task (approaching reward sites vs. reward receipt and consumption). The observed gamma power gradient was highly consistent across all these conditions for both low- and high-gamma (Figure 3A). This systematic power gradient was further confirmed by plotting gamma power as a function of distance from the ventrolateral pole of the probe confirmed the systematic power gradient (Figure 4A and B).

**Figure 3:**
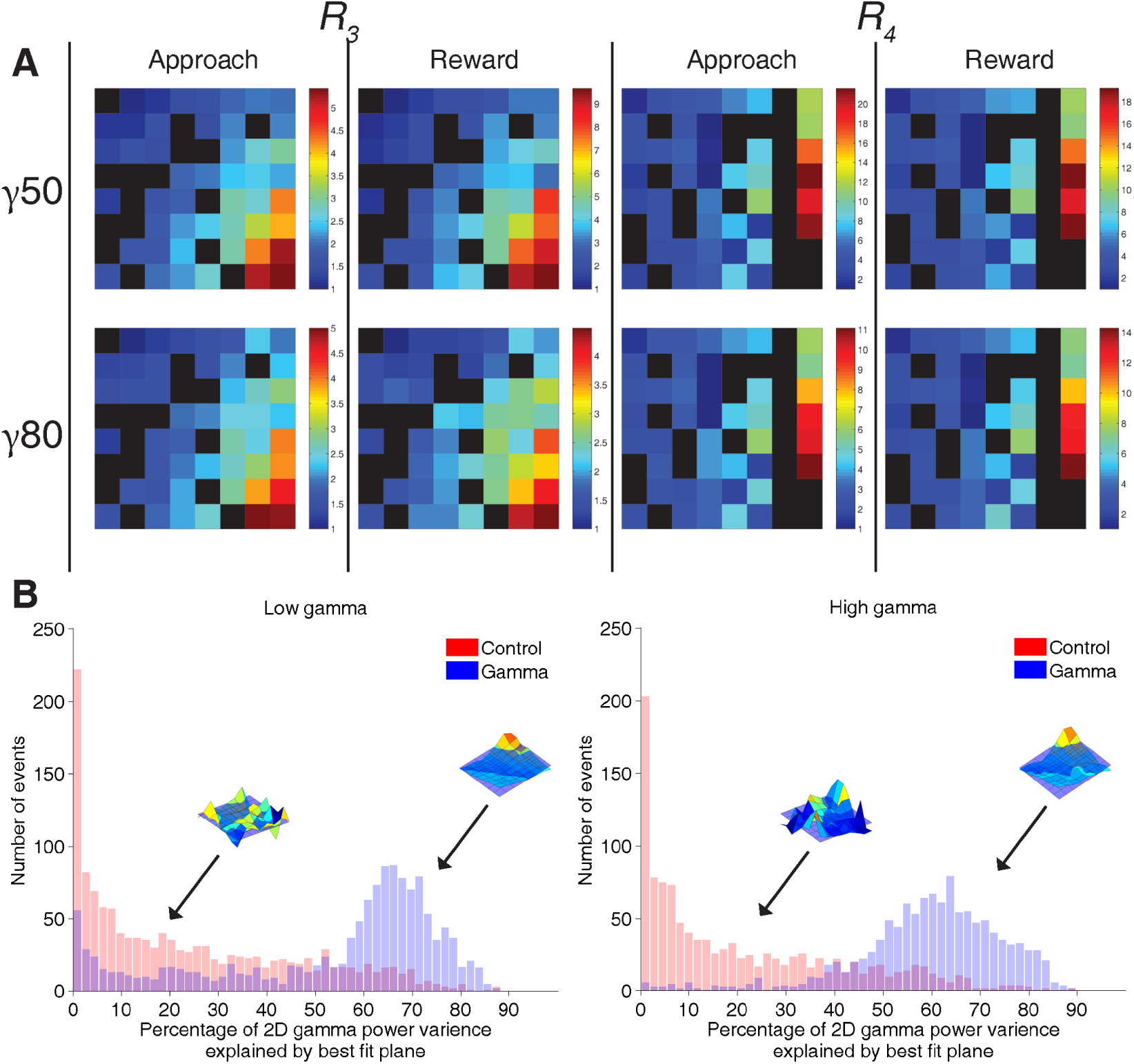
The vStr dorsomedial to ventrolateral gamma power gradient is conserved across different behaviors and is significantly different from randomly selected epochs. **A**: Average gamma power distributions across vStr during active foraging and reward consumption for two subjects that completed the foraging task. **B**: Histograms of variance explained (R^2^) for plane fits to low-gamma (left) and high-gamma (right) and shuffled events of equivalent length (red). Gamma event R^2^ values form a tight cluster with the majority of the variance explained by the best fit plane, while random events fail to fit the plane. Insets are representative gamma and shuffled events.

**Figure 4:**
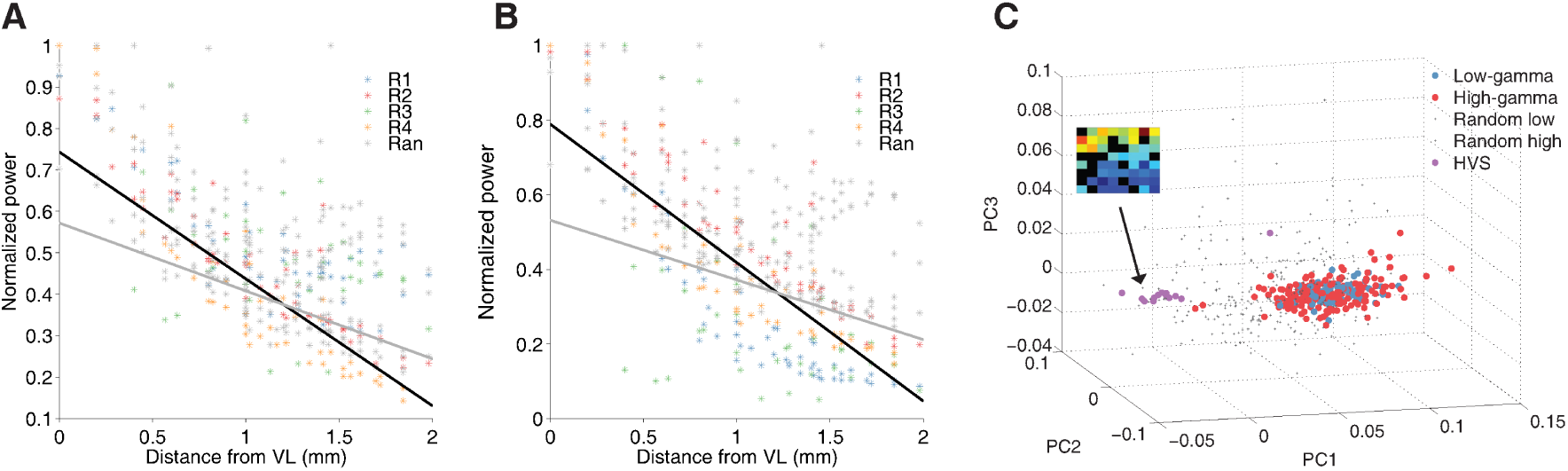
Gamma events form a consistent dorsomedial to ventrolateral power gradient that can be separated from high voltage spindles and random epochs. Random epochs were duration matched periods of low gamma power (see *Methods* for details). **A**: Gamma power relative to distance from the ventrolateral most site on the probe. Average gamma power from each recording session (points) separated by subject (colors) normalized to the maximum value for both low (**A**) and high (**B**). **C**: PCA on the power gradients across events reveals that both low-gamma and high-gamma are indistinguishable from each other, yet are clearly separate from both random epochs and high-voltage-spindles.

To determine if this gradient was simply a consequence of probe impedance values or other peculiarities of the recording setup, we compared the power gradient obtained during detected gamma oscillation events to the power gradient from a set of randomly chosen control events. By fitting a plane to the distribution of power across the probe, we could quantify how much of the variance in power across probe sites was accounted for by a linear power gradient. If this gradient was due to nonspecific properties of the signal, then this pattern would be similar during true gamma oscillations and non-gamma events. Contrary to this scenario, the ventrolateral gradient disappeared for the random control events (Figure 3B, low-gamma events mean R^2^ 52.74 ± 24.08, low-gamma matched random epochs mean R^2^ 22.79 ± 20.94; high-gamma events mean 59.15 ± 15.71, high-gamma matched random epochs mean 18.51 ± 18.11; independent samples t-test: t_(2690)_ = 34.43, p < 0.001 for low-gamma; t_(2308)_ = 57.62, p < 0.001 for high-gamma). Furthermore, as reported previously (Berke et al., 2004), high-voltage spindles (HVS) displayed a power gradient in the opposite direction, with largest power at the dorsomedial pole (Figure 4C inset). Thus, the ventrolateral power gradient observed during gamma oscillations does not result from nonspecific probe or recording system properties.

The ventrolateral gamma power gradient identified above suggests the lack of a local source (in the vStr) for these oscillations. To determine the source of vStr gamma, we first plotted phase differences relative to the ventrolateral recording site. Although the specific pattern of these phase differences varied across subjects, these differences were consistently small (<10^°^, Figure 5A); in particular, there was no evidence of phase reversals, a tell-tale sign of systematically arranged sink/source pairs. The patterns of phase differences were consistent between low- and high-gamma within subjects. Next, we applied current source density (CSD) analysis, which in accordance with the near-zero phase gradients showed only very small sink/source pairs, either in single examples (Figure 6A and B) or when averaged across all events (Figure 6C). Thus no it appears that no obvious source of either low- or high-gamma oscillations exists within the vStr.

**Figure 5:**
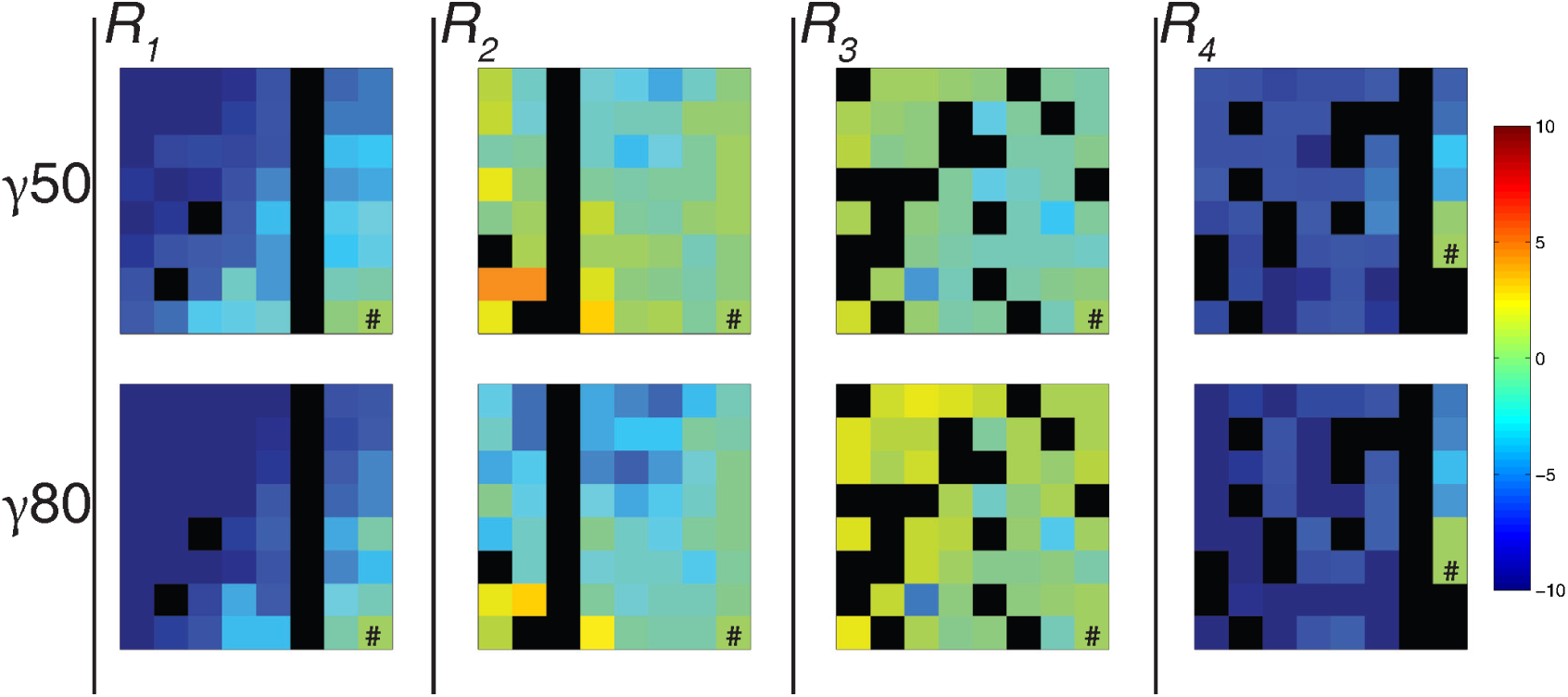
Average phase differences across the center three cycles in each event. Phase lags were found to have a gradient along the ventrolateral to dorsomedial poles, but lacked directional consistency. Each plot shows the average phase difference (in degrees) relative to the ventrolateral most electrode (#). Phase differences were negligible showing a small lag from ventrolateral to dorsomedial in 2/4 subjects. Heat maps range between −10 to 10 degrees (± 0.6ms)

**Figure 6:**
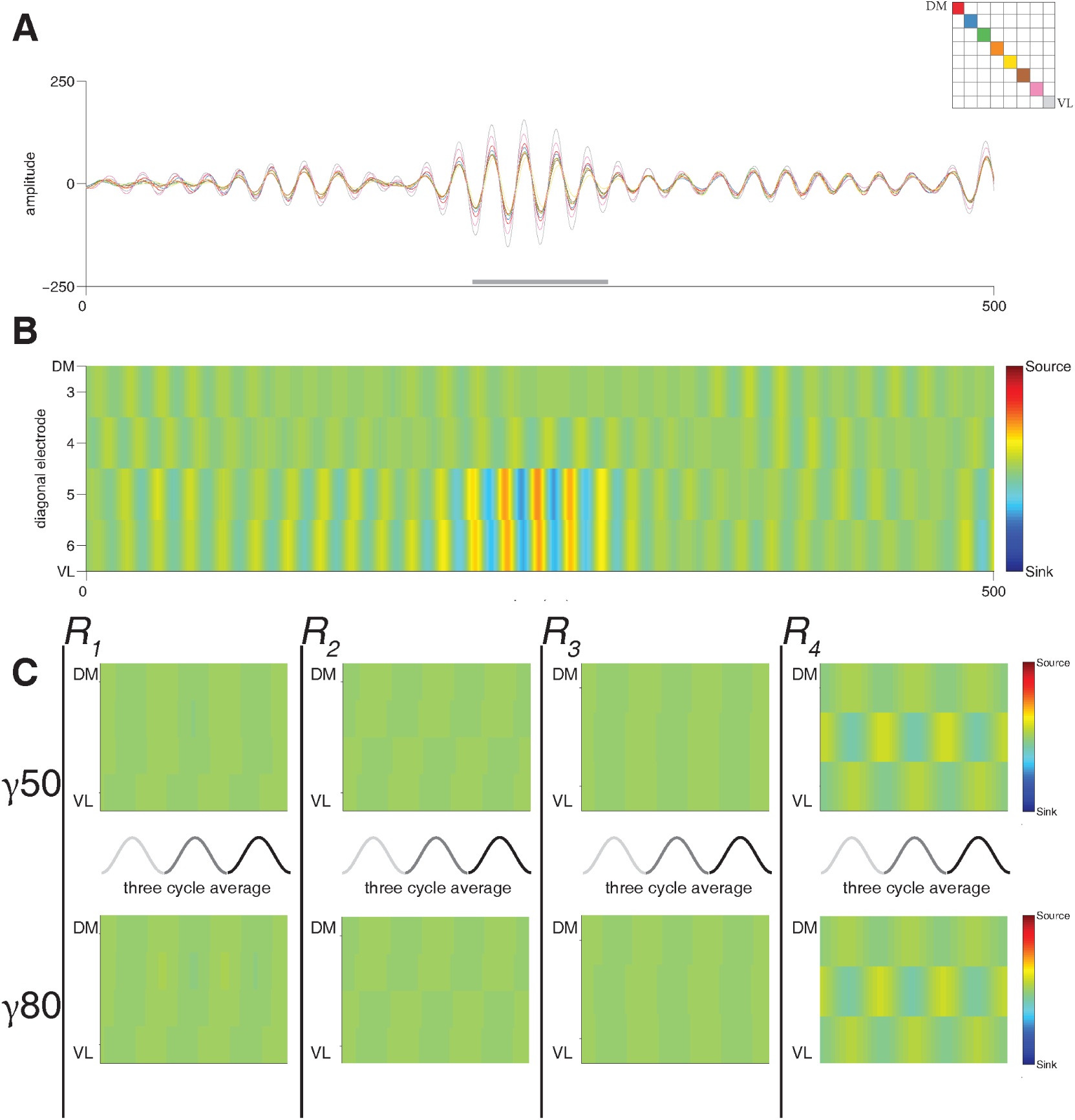
Current source density (CSD) analysis of gamma events. **A**: Filtered low-gamma event (same as shown in Figure 2B). Each trace represents a recording site on the diagonal of the silicon probe (inset), note the change in power along the dorsomedial to ventrolateral axis but very similar phases. **B**: Sample CSD over the same low-gamma event. Pseudocolor scale represents fractional values relative to a 180^°^ phase inversion (source/sink pair). Only a weak source/sink appears on the ventrolateral pole across electrodes, corresponding to the slight phase shift in the example traces. **C**: Average CSD across the center three cycles (grey lines) by subject event. Note that no clear source/sink pair emerges, consistent with the lack of a phase reversal in the gamma-band LFP

### Unilateral naris occlusion strongly reduces vStr gamma power

The clear gradient in vStr gamma power is consistent with Berke (2009)’s proposal that the adjacent piriform cortex, known to generate strong gamma oscillations, may be the main source of vStr LFP gamma. Given that piriform gamma is known to be abolished by occlusion of the ipsilateral nostril (naris, Zibrowski and Vanderwolf 1997) we tested the effects of ipsilateral naris occlusions on vStr gamma power by inserting removable nose plugs alternately in one nostril, and then the other (*Methods*). Contralateral occlusions, performed alternately before or after the ipsilateral condition, provided a control for nonspecific (e.g. behavioral) effects of naris blockage. Ipsilateral naris occlusions effectively abolished low- and high-gamma power relative to the contralateral condition, and relative to unoccluded conditions before (“pre”) or after (“post”, Figure 7A-C and H). Although power spectral densities of individual recording sessions varied, likely due to behavioral differences such as mobility on the pot, gamma suppression was highly consistent across sessions and subjects (Figure 7A-C).

quantify this effect, we compared gamma power extracted from the power spectral density during ipsilateral and contralateral occlusions respectively, after normalizing to gamma power during non-occluded control conditions (“pre” and “post” recording epochs). Only ipsilateral occlusion significantly reduced gamma power in both low- and high-gamma bands to a mean of 0.48 (SEM ± 0.05) and 0.62 (± 0.04) of the control condition. Paired t-tests confirmed the reduction was indeed significant for both low-gamma (t_(11)_ = −10.64, p <0.001) and high-gamma (t_(11)_ = −5.10, p <0.001) relative to the power during the contralateral occlusion. The contralateral occlusion failed to reduce low-gamma power compared to the control (1.00 ± 0.06 of the control condition, t_(11)_ = 0.02, p = 0.99). Contralateral occlusion did produce a marked decrease in the high gamma power relative to the control (0.87 ± 0.05 of the control condition, t_(11)_ = −2.66, p = 0.02). Ipsilateral occlusion provided a significantly greater reduction in gamma power compared to contralateral occlusion for both low- (t_(11)_ = −11.62, p <0.001) and high-gamma (t_(11)_ = −10.79, p <0.001).

This strong reduction in gamma power could be due to fewer gamma events occurring, and/or events having lower gamma power. Gamma event detection applied to the occlusion conditions yielded a significantly lower number of gamma events in the ipsilateral occlusion recording (pre- and post session event count normalized to 1; low-gamma events, 0.04 (± 0.21); high-gamma: 0.13 (± 0.03) compared to the contralateral occlusion (low-gamma: 1.09 ± 0.19, paired t-test t_(11)_ = −5.73, p <0.001; high-gamma: 0.77 ± 0.17, t_(11)_ = −4.10, p <0.005, Figure 7D). The number of events in the ipsilateral occlusion was significantly lower than number of detected events in the unoccluded condition for low-gamma (t_(11)_ = −44.93, p<0.001) and high-gamma (t_(11)_ = −29.59, p <0.001). The contralateral occlusion did not differ compared to the control for low-gamma events (t_(11)_ = 0.47, p = 0.69), or high-gamma events (t_(11)_ = −1.41, p = 0.19). Further supporting the robustness of this result, the ipsilateral condition gamma power and gamma event count was lower than the contralateral condition gamma power in every individual session, without exception. Thus, ipsilateral naris occlusion resulted in a strong reduction in gamma power which resulted in a reduction in the number of events detected.

**Figure 7:**
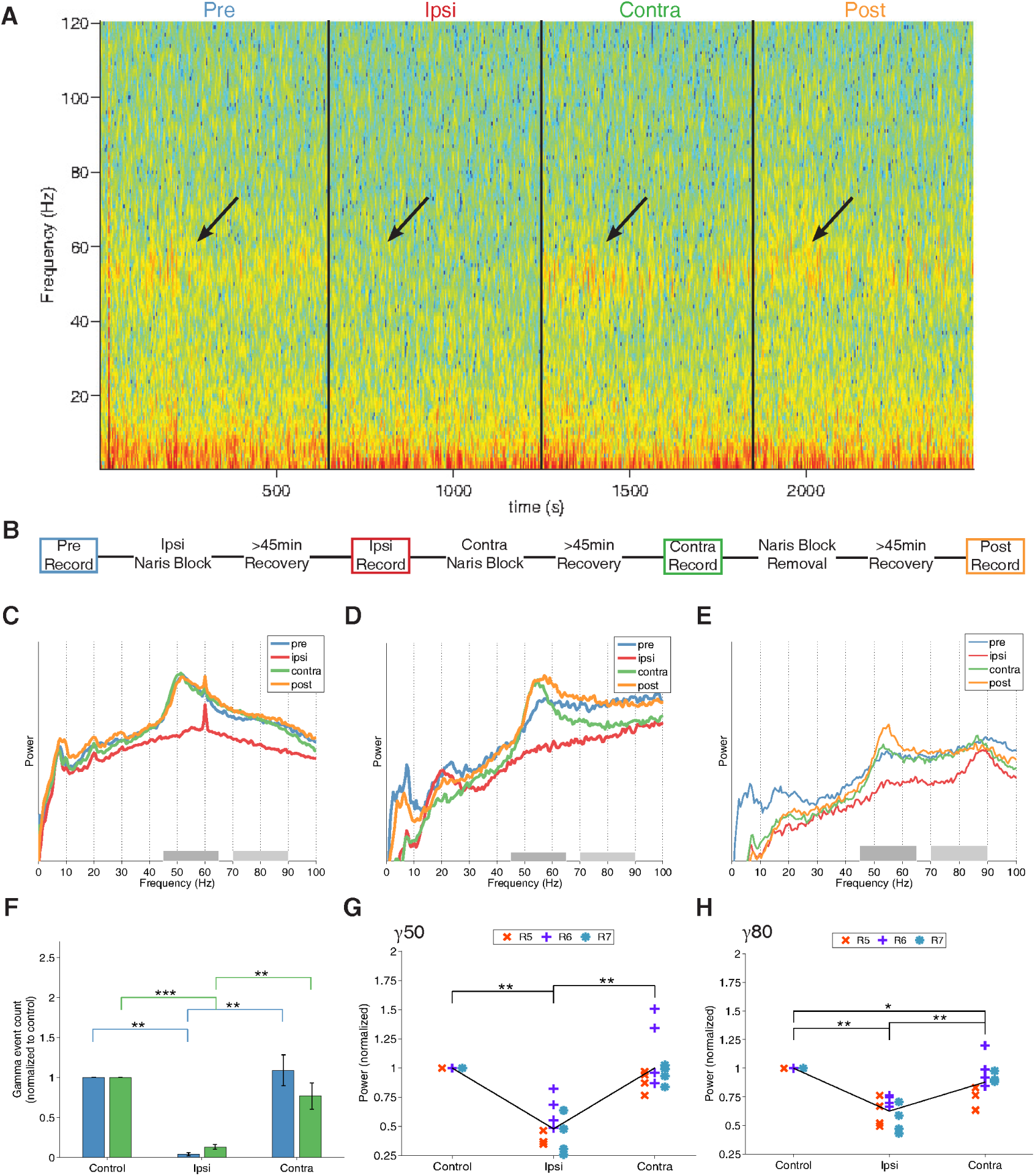
Ipsilateral, but not contralateral, naris occlusion reduces gamma power and event occurrence in the vStr. **A**: Spectrogram across all four experimental phases in a single session. Arrows emphasize the clear gamma band power that disappears during the ipsilateral phase. **B**: Naris experiment timeline. Ipsi- and contralateral occlusion order was counterbalanced across days. **C-E**: Normalized power spectral densities (PSDs) of representative sessions from each rat (R5,6, and 7 respectively). Each session shows a clear reduction in power within the gamma bands for the ipsilateral occlusion condition only (red line). Note that although PSDs differed between sessions (e.g. high-voltage spindles, 7-11 Hz, in the “post” condition in (**D**), the reduction in gamma power was highly consistent. PSDs were computed on the first order derivative of the data to remove the 1/f distributions. **F**: Comparison of the average number of detected gamma events per condition normalized to the unoccluded condition. The ipsilateral condition yielded significantly fewer events for the same recording duration (see main text). Contralateral occlusion increased the number of high-gamma events. Errorbars represent SEM. **G-H**: Comparison of the average power in each session/subject (R4-7) within the low-gamma/high-gamma band. Ipsi- and contralateral conditions were normalized to the unoccluded condition (average between pre and post).

## Discussion

We have demonstrated that (1) the power of gamma oscillations in the ventral striatal local field potential increases along a clear dorsomedial-to-ventrolateral gradient, (2) the phases of gamma oscillations across the vStr are highly consistent, with no evidence of reversals indicating a local sink/source pair, and (3) gamma oscillations were strongly reduced by occlusion of the unilateral, but not contralateral, nostril. Together, these results strongly suggest that gamma oscillations in the vStr LFP are volume-conducted from piriform cortex, consistent with initial observations by Berke (2009), who reported highly similar LFPs in vStr and piriform. Here, we build on this initial work by providing systematic coverage of the vStr with high-density silicon probes, separately analyzing the low- and high-gamma bands and different behaviors, as well as providing a causal manipulation known to disrupt piriform gamma oscillations.

Establishing the source of vStr gamma oscillations is important for at least two distinct reasons. First, knowledge of the source directly informs the interpretation of recorded signals. Because vStr neurons can phase lock to gamma oscillations (Berke, 2009; Kalenscher et al., 2010; van der Meer et al., 2010; Howe et al., 2011; Dejean et al., 2016) and vStr ensemble spiking can predict gamma oscillation frequency (Catanese et al., 2016), the vStr LFP clearly contains at least some information about local (spiking) activity. However, if vStr LFPs contain a component volume-conducted from the piriform cortex, as we have shown, then some changes in the LFP may reflect processing in piriform cortex, rather than local processing in vStr. We expand on the issue of how to reconcile LFP volume conduction with local phase locking below.

The second reason it is important to identify the source(s) of the vStr LFP relates to how that signal is controlled. Several studies have linked properties of the vStr LFP to different task components such as reward approach, receipt, and feedback processing (van der Meer and Redish, 2009; Cohen et al., 2009b), to trait-level variables such as impulsivity (Donnelly et al., 2014), translationally relevant interventions such as manipulations of the dopamine system (Berke, 2009; Lemaire et al., 2012; Morra et al., 2012) and deep brain stimulation (McCracken and Grace, 2009; Doucette et al., 2015). Our results imply that in order to change the vStr LFP, either endogenously or using experimental manipulations, it may paradoxically be more effective to target piriform cortex rather than the vStr itself.

The result that gamma-band LFP oscillations in vStr are primarily volume-conducted from a different structure is in line with the biophysics of LFP generation, which generally require sink/source pairs of transmembrane currents to be aligned so that their contributions may sum spatially to generate systematic changes in the LFP (Nunez and Srinivasan, 2006; Buzsáki and Wang, 2012). The striatum, as a non-layered structure with generally radially symmetric dendritic arbors (Kawaguchi et al. 1995;Tepper et al. 2004, 2010), lacks the organization conducive to spatial summing of currents. Nevertheless, there have been reports of local heterogeneity in vStr gamma oscillations. For instance, Kalenscher et al. (2010) and Morra et al. (2012) show example recordings for which specific channels show phases or amplitudes apparently inconsistent with volume conduction. When we found such examples in our data, however, they could be attributed to impedance magnitude or angle changes on isolated electrode sites; the high-density view afforded by Si probe recordings can disambiguate these cases. A different body of work has suggested that high-frequency oscillations can be generated locally in the vStr because they are affected by infusions of MK801 and lidocaine into the vStr (Hunt et al., 2010; Olszewski et al., 2013); however, given its anatomical proximity, it is conceivable that some of the drugs spread to act on the piriform cortex.

So, what is the correct interpretation of ventral striatal gamma oscillations in the LFP, given the apparent paradox of evidence for volume conduction on the one hand (as presented here, stylized in Figure 8A) and evidence for local phase locking and ensemble coding to gamma-band LFP on the other? We found no phase reversals that would indicate local generation of gamma oscillations in the LFP (Figure 8B), as could be supported by cortical pyramidal-interneuron circuits as demonstrated in neocortical areas (Cardin et al., 2009; Sohal et al., 2009; Siegle et al., 2014), or by cell-intrinsic resonance (Taverna et al., 2007). A different possibility is that LFP oscillations result from local transmembrane currents, but are inherited from inputs to the vStr through the synaptic currents they generate (Figure 8A). As with the local generation scenario, our results seem to rule out this possibility, given that the distribution of gamma power across the vStr does not appear to match known anatomical distributions (Figure 8D; reviewed in Groenewegen et al. 1999; Humphries and Prescott 2010), and disappears with piriform inactivation. The vStr does receive inputs from piriform cortex, however (Brog et al., 1993; Schwabe et al., 2004), which could account for phase-locking in the vStr: to the extent that piriform cortex inputs are effective in driving vStr spiking, then that spiking would be expected to lock to the field potential originating in the same source structure (Figure 8C).

**Figure 8:**
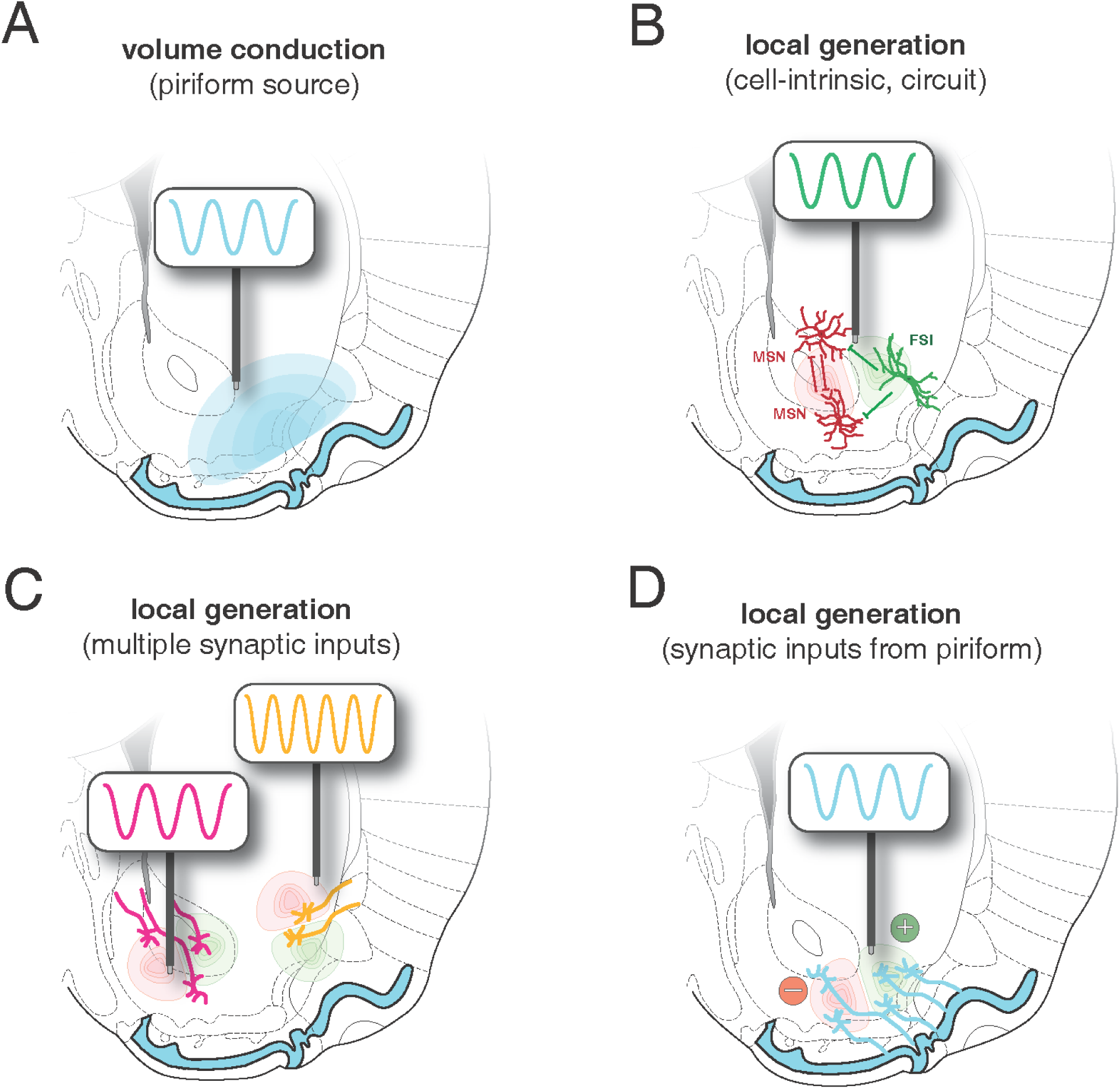
Four possible scenarios for the generation of gamma-band oscillations in the vStr. **A**: Volume conduction originating in the adjacent piriform cortex would show no clear phase reversals within the vStr. **B**: Local mechanisms such as cell-intrinsic currents and circuitry produce local gamma oscillations within the vStr circuit. **C**: Local sources matching the anatomical heterogeneity of the vStr, here idealized with two different afferent sources (orange and magenta). **D**: Rhythmic inputs from the adjacent piriform cortex lead to local generation within the vStr, which should follow anatomical projection densities. Our data did not show phase reversals within the vStr ruling out local generation by cell-intrinsic or multiple synaptic inputs. Inactivation of the piriform cortex greatly reduced gamma oscillations across the vStr making volume conduction the most plausible source of vStr gamma oscillations in the LFP, but does not rule out inherited inputs from the piriform, though a lack of phase reversals makes this unlikely.

The above interpretation of vStr gamma oscillations has implications for a number of avenues of research involving the vStr. For instance, a long-standing notion is that vStr may provide a “switchboard” between inputs from prefrontal cortex, amygdala, and hippocampus (O’Donnell and Grace, 1995; Gruber et al., 2009); LFP oscillations are a major candidate for implementing and/or reflecting such functions (Fries, 2005, 2015). Our results suggest that vStr circuits are unlikely to contain the “controls” that determine the timing and frequency of vStr gamma oscillations. Instead, gamma oscillations volume-conducted and/or inherited from a common piriform source may be a powerful synchronizing drive of neural activity in the rodent limbic system. In particular, LFP synchrony in the gamma band across limbic structures such as prefrontal cortex, orbitofrontal cortex, ventral hippocampus, and amygdala (van Wingerden et al., 2014; Harris and Gordon, 2015; Catanese et al., 2016), may, at least in rodents, be shaped by piriform input. Given the much larger distance from the human vStr to piriform cortex, and the widespread use of relatively local referencing in depth electrode recordings, it seems a priori unlikely that gamma oscillations in the human vStr are volume conducted from piriform cortex. More generally, however, there is at least some evidence that lateralized nasal breathing affects both the EEG signal and various aspects of cognitive performance (Block et al., 1989; Zelano et al., 2016); intracranial EEG recordings in epilepsy patients show a connection between nasal breathing and increases in power of human delta (0.5-4 Hz), theta (4-8 Hz), and beta (13-34 Hz) oscillations in the piriform, amygdala and hippocampus. Although gamma activity has been linked to respiration in the olfactory circuit in rodents (Gault and Leaton, 1963), the time course of vStr gamma power rules out respiration as the only factor controlling gamma oscillations in the vStr LFP. For instance, several studies have noted strong suppression of gamma power as animals run, compared to rest (van der Meer and Redish, 2009; Malhotra et al., 2015).

Our study has a number of limitations: we chose to focus on gamma-band oscillations for several reasons, including the high consistency with which these oscillations can be probed across multiple species (rodents and humans in particular), because it is the vStr oscillation band which has received the most attention in terms of behavioral correlates and relationship to spiking activity, and because gamma oscillations are plentiful during rest and well as during behavior. However, clearly it would be of interest to determine the sources of other oscillations in the vStr LFP, such as delta, theta and beta, which have all been linked to local spiking activity and behavior (Van der Meer and Redish, 2011; Howe et al., 2011; Stenner et al., 2015; Malhotra et al., 2015). The data we recorded as part of this study did not reliably contain clearly identifiable epochs with these oscillations, so this is an avenue for further work. Also, our naris occlusion procedure likely affects olfactory areas in addition to piriform cortex, such as the olfactory bulb and the olfactory tubercle (in rats: Zibrowski and Vanderwolf 1997); however, owing to its large size, convoluted shape and positioning at the ventral surface of the brain, piriform cortex is difficult to target with higher specificity. Centrifugal afferents from the entorhinal and piriform cortices and olfactory tubercle are capable of modulating olfactory bulb gamma even in the absence of the main peduncle input (Gray and Skinner, 1988), yet we see a strong suppression of gamma power, suggesting that the entire circuit is sufficiently impaired. Despite this limitation, we point to the convergence between the naris occlusion experiment and the power gradient observed in the probe recordings to support the most parsimonious interpretation that gamma LFP oscillations in the vStr originate in piriform cortex.

In closing, we wish to stress an important point: the above conclusion that vStr gamma LFP oscillations are volume-conducted from piriform cortex does *not* mean oscillations in the vStr LFP are not important or an epiphenomenon. As pointed out earlier, the spiking of vStr neurons shows clear oscillatory signatures, including intrinsically generated resonance in the gamma range (Taverna et al., 2007). Indeed, given that pretty much any input to the vStr is known to have oscillatory activity at the LFP and spiking levels, it would be hard to imagine how vStr activity would not itself also show oscillations, which in turn can be used as an access point to define and manipulate specific functional sub-populations and state changes, as has been tremendously successful in other areas (Pesaran et al. 2002; Colgin et al. 2009; Bosman et al. 2012). Our results should motivate care in the interpretation of the vStr LFP, and suggest future work in determining how olfactory inputs may shape activity not just in the vStr but other limbic structures.

## Acknowledgments

We thank Nancy Gibson, Martin Ryan and Jean Flanagan for animal care, Claire Cheetham for suggesting the naris occlusion technique, Youki Tanaka for reagents, and Min-Ching Kuo and Alyssa Carey for technical assistance. This work was supported by Dartmouth College (Dartmouth Fellowship to JEC, and start-up funds to MvdM) and the Natural Sciences and Engineering Research Council (NSERC) of Canada (Discovery Grant award to MvdM, Canada Graduate Scholarship to JMG).

